# Where there’s smoke, there’s fuel: dynamic vegetation data improve predictions of wildfire hazard in the Great Basin

**DOI:** 10.1101/2021.06.25.449963

**Authors:** Joseph T. Smith, Brady W. Allred, Chad S. Boyd, Kirk W. Davies, Matthew O. Jones, Andrew R. Kleinhesselink, Jeremy D. Maestas, David E. Naugle

## Abstract

Wildfires are a growing management concern in western US rangelands, where invasive annual grasses have altered fire regimes and contributed to an increased incidence of catastrophic large wildfires. Fire activity in arid, non-forested ecosystems is thought to be largely controlled by interannual variation in fuel amount, which in turn is controlled by antecedent weather. Thus, long-range forecasting of fire activity in rangelands should be feasible given annual estimates of fuel quantity. Using a 32 yr time series of spatial data, we employed machine learning algorithms to predict the relative probability of large (>405 ha) wildfire in the Great Basin based on fine-scale annual and 16-day estimates of cover and production of vegetation functional groups, weather, and multitemporal scale drought indices. We evaluated the predictive utility of these models with a leave-one-year-out cross-validation, building spatial hindcasts of fire probability for each year that we compared against actual footprints of large wildfires. Herbaceous aboveground biomass production, bare ground cover, and long-term drought indices were the most important predictors of burning. Across 32 fire seasons, 88% of the area burned in large wildfires coincided with the upper 3 deciles of predicted fire probabilities. At the scale of the Great Basin, several metrics of fire activity were moderately to strongly correlated with average fire probability, including total area burned in large wildfires, number of large wildfires, and maximum fire size. Our findings show that recent years of exceptional fire activity in the Great Basin were predictable based on antecedent weather-driven growth of fine fuels and reveal a significant increasing trend in fire probability over the last three decades driven by widespread changes in fine fuel characteristics.

## Introduction

Shifting fire regimes in the western US present escalating management challenges. Though a century of aggressive fire suppression has resulted in an overall deficit of fire (Marlon et al. 2012), the annual area burned and incidence of large wildfires (>405 ha) have risen in recent decades (Dennison et al. 2014, Westerling 2016, Iglesias et al. 2022). Humans have increased ignitions (Balch et al. 2017), altered fuel characteristics (Noss et al. 2006, Balch et al. 2013, Fusco et al. 2019), and otherwise modified fire regimes of many western ecosystems, with impacts including permanent vegetation state transitions (D’Antonio and Vitousek 1992, Coop et al. 2020) and degradation or loss of habitat for sensitive species (Rockweit et al. 2017, O’Neil et al. 2020). Responding to these trends and threats, the US federal fire suppression budget has grown more than six-fold in the past three decades, from an average of $370M·yr^-1^ from 1985–1989 to $2.38B·yr^-1^ from 2016–2020 (National Interagency Fire Center, 2021). Even so, widespread and synchronous fire activity across western North America increasingly exceeds fire suppression capacity (Podur and Wotton 2010, Abatzoglou et al. 2021), contributing to increased incidence of wildfire disasters and underscoring the need for accurate, long-range forecasts to guide management and allocation of fire suppression resources.

Although forest fires have historically been the focal point of fire management and research, increasing wildfire in non-forested ecosystems across western North America (hereafter rangelands) is a growing concern. The Great Basin, a cold desert covering most of Nevada and parts of California, Idaho, Oregon, and Utah, has recently experienced particularly damaging fire seasons fueled in part by highly flammable exotic annual grasses such as cheatgrass (*Bromus tectorum* L.; Balch et al. 2013, Fusco et al. 2019). Native vegetation communities of the Great Basin, co-dominated by perennial grasses and non-resprouting shrubs such as sagebrush (*Artemisia* spp.), evolved with infrequent fire and can take many decades to recover after burning (Knutson et al. 2014, Shriver et al. 2018, Bates et al. 2020). Larger and more frequent wildfires therefore threaten to catalyze widespread, permanent shifts to vegetation communities devoid of shrubs and dominated by exotic annual grasses, with negative consequences for rural communities and shrubland obligate species (Knick et al. 2003, Coates et al. 2016, O’Neil et al. 2020).

Spatial tools quantifying wildfire hazard fall into two broad categories: static land cover- and climate-based maps depicting the long-term average annual probability of burning (e.g., Short et al. 2020), and short-term (daily to weekly) indices based on rapidly changing variables such as fuel moisture, humidity, wind speed, and temperature (e.g., the Keetch-Byram Drought Index [Keetch and Byram 1968] and the various indices under the National Fire Danger Rating System [Deeming et al. 1972, Burgan et al. 1998]). These tools reflect a historical focus on “flammability-limited” forested ecosystems, where fuel quantity changes little from one year to the next but fire danger can change rapidly within a season as the moisture content of fuels responds to atmospheric conditions (Krawchuk and Moritz 2011, Abatzoglou and Kolden 2013).

Semi-arid and arid grasslands and shrublands, in contrast, are considered “fuel-limited” in the sense that burnable biomass can vary considerably from year to year while fuel moisture is rarely limiting (Krawchuk and Moritz 2011, Abatzoglou and Kolden 2013). In these ecosystems, fire activity is strongly correlated with antecedent (i.e., prior to the onset of the fire season) precipitation and fuel growth, and relatively weakly related to weather during the fire season (Westerling et al. 2002, 2003, Brown et al. 2005, Littell et al. 2009, Abatzoglou et al. 2018). Consequently, fire activity at the seasonal scale may be predictable further in advance in rangelands than in forested ecosystems.

Precipitation and drought indices have long been employed as correlates of fine fuel growth for predicting fire activity in rangelands. Outlooks distributed months in advance of the fire season, for example, rely heavily on long-term drought indices (e.g., Palmer Drought Severity Index) to infer the relative quantity of fine fuels. Recent work has strengthened and refined the links between precipitation and fine fuels in Great Basin shrublands. Pilliod et al. (2017) found precipitation had complex, multi-year lagged effects on fuel accumulation that differed among plant functional groups and concluded that native perennial bunchgrass production in the previous year, as well as litter accumulated over 1–3 years’ production of exotic grasses and forbs, were among the main drivers of the number of large fires and total burned area in the Great Basin. Using only antecedent precipitation variables, they developed annual fire risk maps predictive of the distribution of large fires across the Great Basin (Pilliod et al. 2017).

Precipitation and drought are, however, coarse proxies for fine fuel accumulation. Direct, remotely-sensed estimates of vegetation cover and/or production may better account for fine-grained variation induced by factors such as topography and soil, invasion by exotic annual grasses, and legacies of management and disturbance. For example, MODIS-derived estimates of production revealed high intra- and interannual variation in fuels driven by the combined effects of weather and disturbance history in northern Great Basin shrublands (Li et al. 2020). Despite its potential utility, the application of remotely sensed data to wildfire forecasting in rangelands has been hampered by a lack of readily available, timely data quantifying dynamic and spatially variable rangeland fuels at ecoregional and larger extents.

Recently, datasets providing high-resolution, consistent, annual estimates of rangeland vegetation cover and production with extensive temporal and geographic coverage have been developed from long-term satellite imagery records (e.g., Jones et al. 2018, Zhang et al. 2019, Rigge et al. 2020). These dynamic vegetation datasets have been rapidly adopted for quantifying long-term vegetation trends (Robinson et al. 2019, Fogarty et al. 2020, Rigge et al. 2021), wildlife habitat (Donovan et al. 2021, Olsen et al. 2021, Pilliod et al. 2022), spread of invasive species (Pastick et al. 2021, Smith et al. 2022), and outcomes of management (Bestelmeyer et al. 2021, Fick et al. 2022). To our knowledge, however, they have yet to be used in a wildland fire preparedness context. Our overarching objective was to explore the utility of these data for quantifying rangeland fuels and improving wildfire preparedness in the Great Basin.

Specifically, we use vegetation cover and production data from the Rangeland Analysis Platform (hereafter RAP), a suite of dynamic rangeland vegetation datasets based on the extensive Landsat imagery record and decades of intensive ground sampling (Jones et al. 2018). RAP provides annual estimates of cover of rangeland plant functional types (annual forbs and grasses, perennial forbs and grasses, shrubs, and trees), litter, and bare ground over the coterminous U.S. from 1984–present at a 30 m resolution (Allred et al. 2021). In addition to cover, RAP provides annual production (kg·ha^−1^) estimates for all plant functional types and 16-day production (kg·ha^−1^·16 d^-1^) estimates for herbaceous plant functional types (Jones et al. 2021).

Our aim was to use vegetation cover and production data from RAP in conjunction with gridded weather, drought, and climate data to produce annual, high-resolution (120 m) forecasts of the relative probability of burning in a large (>405 ha) wildfire available well before the onset of the fire season and to quantify the skill of these forecasts using a hindcasting (i.e., forecasts for past years) approach. Numerous factors, many unrelated to fuels and subject to change over time (e.g., ignition probability), influence where and when large wildfires occur (Balch et al. 2018). Our objective was not to account for all these factors, but rather to develop a fuels-based index proportional to the annual probability of burning. An additional objective was to explore the trade-off between forecast lead-time and accuracy. We anticipated that short-term drought indices and herbaceous vegetation production in the spring would become increasingly informative as the forecast date approached the onset of the fire season, and therefore forecasts made earlier in the year, when they would potentially be more useful for planning, would be less accurate than forecasts made later in the spring. Finally, to explore the utility of our approach relative to currently available spatial fire risk products we compare the skill of our hindcasts to a static burn probability map widely used by land management agencies (Short et al. 2020) in predicting the spatial patterns of large wildfires over the last 3 decades.

## Methods

### Study area and period

We focused our analysis on rangelands as defined by Reeves and Mitchell (2011) in the Great Basin. EPA Level III ecoregions (Omernik and Griffith 2014) defined the boundaries of our study area, which included the Central Basin and Range, Northern Basin and Range, and Snake River Plain ecoregions. Data availability constrained our analysis to fire seasons from 1988 to 2019.

### Data sources

Annual estimates of vegetation cover (percent) and aboveground biomass production (kg·ha^−1^), and short-term estimates of herbaceous aboveground biomass production (kg·ha^−1^·16-d^-1^), were derived from the Rangeland Analysis Platform (RAP; Jones et al. 2018, 2021, Allred et al. 2021). Annual cover and production, which are updated each year in early spring with estimates for the previous year’s growing season, were included at 1- and 2-yr lags; i.e., we used cover and production in year *t*-2 and *t*-1 to predict the occurrence of fire in year *t*. The short-term (16 d) herbaceous production dataset is updated in near-real-time as provisional Landsat images are made available, and was used to quantify cumulative growth of fine fuels in year *t* from January 1 until the date the forecast was made.

Water availability is a dominant control on primary production in most terrestrial ecosystems (Churkina and Running 1998), and precipitation is the primary driver of interannual variation in herbaceous vegetation cover in Great Basin rangelands (Pilliod et al. 2017). We therefore included precipitation and drought variables derived from the Gridded Surface Meteorological dataset (gridMET; Abatzoglou 2013) as indicators of resource availability for vegetation growth at multiple temporal scales. Precipitation was quantified over the preceding year (pr_annual) and over the water year-to-date (pr_wytd; October 1 of the previous year through the forecast date). Although the Palmer Drought Severity Index (PDSI; Palmer 1965) has historically seen wide use in fire forecasting (Westerling et al. 2002, 2003), newer drought indices have been developed that allow drought to be quantified over various user-specified temporal scales. We explored two of these indices, the standardized precipitation-evapotranspiration index (SPEI; Vicente-Serrano et al. 2010) and evaporative demand drought index (EDDI; Hobbins et al. 2016, McEvoy et al. 2016), calculated at 30 d, 90 d, 180 d, 270 d, 1 yr, 2 yr, and 5 yr temporal scales. The SPEI and EDDI both range from approximately -2 to 2, but their interpretations are opposite: negative values of SPEI indicate drought, whereas negative values of EDDI indicate wetter conditions (less evaporative demand). These indices are updated every 5 days and are available for the period 1980–present.

Low temperature limits vegetation growth during a significant portion of the year in the Great Basin. With adequate moisture, more rapid spring warming should facilitate faster vegetation growth and greater accumulation of fine fuels. We therefore calculated accumulated growing degree days (gdd) from January 1 to the date of the forecast using daily minimum and maximum temperatures from gridMET.

Climate may directly or indirectly influence the annual probability of burning. A direct effect might include variation in the annual number of days with weather conducive to wildfire ignition and spread. Climate may also indirectly affect annual probability of fire, e.g. via vegetation community composition. Vegetation communities occupying different biophysical settings are likely to differ in their responses to precipitation, temperature, and drought. To account for this spatial climate variation, we included mean annual temperature and mean annual precipitation from the PRISM 30-yr (1991–2020) normal dataset (Daly et al. 2015).

We derived the categorical response, burned (*y* = 1) or unburned (*y* = 0) from the Monitoring Trends in Burn Severity dataset (MTBS; Eidenshink et al. 2007). MTBS maps all fires >405 ha (>1000 ac) since 1984 across all land ownerships. Unburned inclusions often occur within MTBS mapped fire perimeters (Kolden et al. 2012), introducing opportunity for class label errors. Therefore, we used the MTBS thematic burn severity raster dataset to assign each pixel to the burned or unburned class. Thematic burn severity classes 1–5 indicate increasing burn severity. We assigned all pixels in burn severity class 1 (low severity/unburned) or outside fire perimeters to the unburned class (*y* = 0), and all pixels in classes 2–5 to the burned class (*y* = 1). Class 6 represents pixels of unknown burn severity within fire perimeters. We assumed pixels in class 6 were assigned this class due to imagery issues (e.g., clouds or shadows), not because they were unburnable land cover types such as large water bodies; unburnable land cover types were already masked from our analysis. Therefore, we assigned pixels in class 6 to the burned class (*y* = 1).

Because MTBS only maps fires >405 ha, some sampled pixels assigned to the unburned class may have, in fact, burned in small wildfires. However, using data from the USFS Fire Program Analysis Fire Occurrence dataset (Short 2021), we estimated that fires <405 ha affected, on average, only ∼0.05% of the study area not burned in MTBS-mapped fires each year. We therefore assumed the level of contamination of the unburned sample was negligible. All data were accessed via Google Earth Engine (GEE; Gorelick et al. 2017), and analyses were conducted using the GEE code editor and R (version 4.0; R Development Core Team, 2021).

### Modeling approach

We used random forests (RF; Breiman 2001) to predict the response variable (burned or unburned). Random forests are widely used for prediction in ecology because they are straightforward to fit, require little tuning, and readily accommodate non-linear relationships and interactions among predictors (Cutler et al. 2007). Because they are based on decision trees, predictive accuracy of RF is robust to collinearity among predictors. Importantly for our application, RF generally perform well at predicting probabilities (Niculescu-Mizil and Caruana 2005).

Like many classification algorithms, however, RF performance is degraded when training data are highly imbalanced among classes (Sage et al. 2020). Because only a small fraction of the study area burns annually, a simple random sample would be dominated by unburned pixels. We therefore stratified by outcome to train the model on equal numbers of burned and unburned pixels. This sampling scheme results in pixel-level model predictions, ranging from 0– 1, that do not represent absolute probabilities of burning. However, our goal was not to estimate absolute probabilities of burning, but rather to quantify the relative probability of burning in a large wildfire based on fuel characteristics. We define this quantity *p* at a given pixel with observed predictors *x, p* = *P*(*y* = 1|*x*). Estimates of probabilities 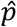 were derived from RF classification models using tree vote aggregation (Sage et al. 2020). Each tree *t* predicts the most probable class, *ŷ*_*t*_ ∈ {0,1} (‘casts a vote’), and 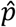 is derived by averaging votes across *T* trees, 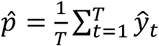.

### Workflow

Our goal was to develop and tune a RF model that can be updated annually to produce a pre-fire season forecast using data sampled from the extensive historical record available in GEE. Although GEE is a powerful platform for accessing, processing, and sampling data, fitting a model, and mapping predictions across large areas, it has limited built-in statistical functions. Therefore, hyperparameter tuning and statistical evaluation of hindcasts were accomplished outside of GEE. The basic outline of our workflow is shown in Fig. 1.

**Figure 1.**
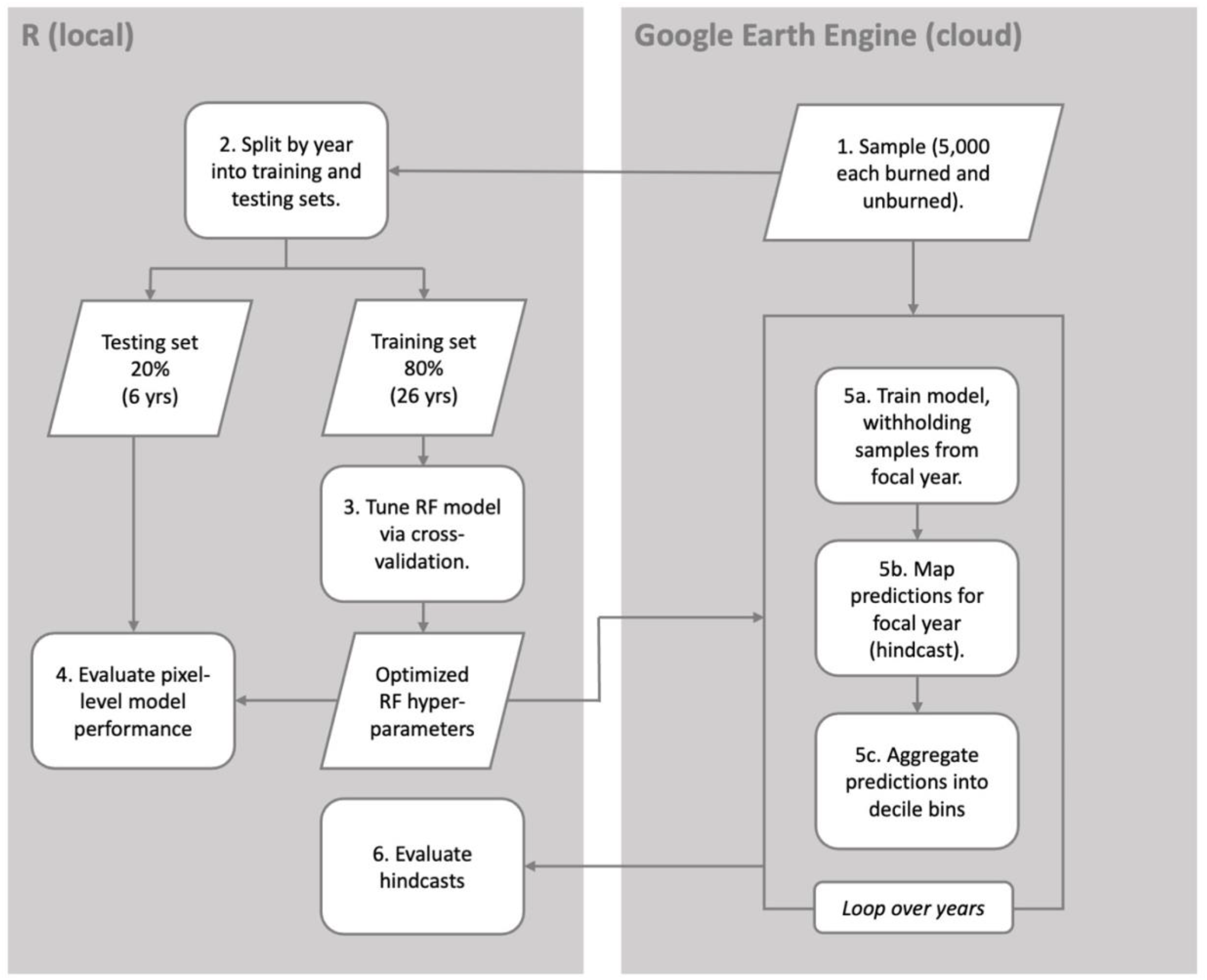
Workflow used to develop and evaluate dynamic spatial wildfire forecasts based on antecedent cover and production of vegetation, multitemporal scale drought indices, and climate variables for the Great Basin, USA.

#### 1. Sampling

We drew a sample of *n* = 10,000 pixels (5,000 each burned and unburned) for model tuning and testing. Native resolution of response and predictor datasets varied from 30 m to 4.6 km (Table 1); we chose to resample all data to a common 120 m resolution for model fitting and prediction. Training data were sampled randomly in time and space to reduce spatial and temporal autocorrelation among training data. This is particularly important for modeling a process like fire; in any given fire season, the outcomes of spatially adjacent pixels are strongly interdependent. Our sampling schema ensured only a single year’s observation, randomly selected from the 32 yr time series, could be included at any given spatial coordinates. Thus, even when sampled pixels were in close spatial proximity, it was unlikely the observations represented by those samples were proximal in time, and vice versa. This sample size was chosen to balance predictive accuracy and processing time; in preliminary analysis, the incremental improvements in accuracy yielded by larger samples did not justify the increase in processing time. This was likely related to the fact that only 2,201 large wildfires overlayed our study area; additional burned samples would largely be drawn from fires already sampled, and therefore yield diminishing returns in terms of new information.

**Table 1.** Predictors used in random forest models of large rangeland wildfire occurrence in the Great Basin, USA, 1988–2019.

| Predictor | Abbreviation | Native resolution<sup>1</sup> | Temporal period/scale | Source<sup>2</sup> |
| --- | --- | --- | --- | --- |
| Annual forb and grass cover | AFGC | 30 m | Annual, lag-1 and lag-2 | 1 |
| Perennial forb and grass cover | PFGC | 30 m | Annual, lag-1 and lag-2 | 1 |
| Shrub cover | SHR | 30 m | Annual, lag-1 | 1 |
| Litter cover | LTR | 30 m | Annual, lag-1 and lag-2 | 1 |
| Bare ground cover | BG | 30 m | Annual, lag-1 and lag-2 | 1 |
| Tree cover | TREE | 30 m | Annual, lag-1 | 1 |
| Annual forb and grass aboveground biomass production | afgAGB | 30 m | Annual, lag-1 and lag-2 | 2 |
| Perennial forb and grass aboveground biomass production | pfgAGB | 30 m | Annual, lag-1 and lag-2 | 2 |
| Herbaceous aboveground biomass production | herbAGB | 30 m | Annual, lag-1 and lag-2 | 2 |
| Shrub net primary production | shrNPP | 30 m | Annual, lag-1 and lag-2 | 2 |
| Cumulative 16-d annual forb and grass aboveground biomass production | afgAGB_ytd | 30 m | Jan 1-forecast DOY | 2 |
| Cumulative 16-d perennial forb and grass aboveground biomass production | pfgAGB_ytd | 30 m | Jan 1-forecast DOY | 2 |
| Standardized Precipitation-Evapotranspiration Index | spei | 4.6 km | 30 d, 90 d, 180 d, 270 d, 1 yr, 2 yr | 3 |
| Evaporative Demand Drought Index | eddi | 4.6 km | 30 d, 90 d, 180 d, 270 d, 1 yr, 2 yr | 3 |
| Precipitation | pr | 4.6 km | Annual (ann) and water year to date (wy) | 4 |
| Cumulative growing degree days | gdd | 4.6 km | Jan 1-forecast DOY | 4 |
| Mean annual temperature | bio01 | 800 m |  | 5 |
| Mean annual precipitation | bio12 | 800 m |  | 5 |

<sup>1</sup>All data were resampled to a common 120 m resolution for model fitting and prediction.
<sup>2</sup>Sources:
- 1: Rangeland Analysis Platform (RAP) fractional cover dataset (Jones et al. 2018, Allred et al. 2021). - 2: RAP annual and 16-day production dataset (Jones et al. 2021) - 3: gridMET Drought Indices; University of California Merced (Abatzoglou 2013). - 4: gridMET; University of California Merced (Abatzoglou 2013) - 5: PRISM 30-yr (1991–2020) climate normals (Daly et al. 2015)

#### 2. Dividing into training and testing data

We split the sample into a tuning set (∼80%) and testing set (∼20%) by randomly selecting 6 of the 32 years and withholding all samples from those years to evaluate the final model.

#### 3. Tuning via cross-validation

RF defaults are to build each tree on a bootstrap sample of the training data (sampling fraction = 1, sampled with replacement) and consider a random selection of 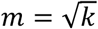 of the *k* available predictors at each split (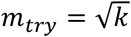; Breiman 2001). We performed a grid search to select the optimal combination of *m*_*try*_ and sampling fraction for our application. We considered values of *m*_*try*_ from 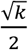 to 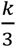, and trees grown on subsamples (without replacement) of the training data ranging in size from 0.1*n* to 0.5*n*, or the default bootstrap sample (with replacement) of size *n* (Martínez-Muñoz and Suárez 2010). We held the minimum node size and maximum number of nodes constant at values (1 and unlimited, respectively) recommended for probability estimation using tree vote aggregation (Liaw and Wiener 2012, Sage et al. 2020). RF models were fit with the `ranger` function in the “ranger” package (version 0.12.1; Wright and Ziegler 2017) in R. All forests were grown to 1,001 trees.

We used cross-validation to assess each combination of tuning parameters. Cross-validation folds were defined using the natural grouping structure of the data; one year’s samples were withheld at a time for prediction. When accurate class probabilities (e.g., relative probability of burning) are the primary objective, the log-loss is more informative than threshold-based metrics such as percent correctly classified, area under the receiver operating characteristic curve (AUC), or kappa. For a predicted fire probability 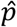 and observed outcome *y*, the log-loss is defined as:

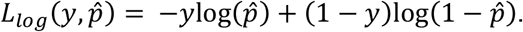

The equation can be modified to place larger penalties on “bad” probabilities (i.e., outcomes predicted to be highly improbable) for a particular class by introducing a weighting parameter, *α*:

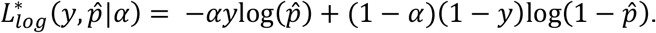

For our purposes, a pixel may remain unburned despite suitable fuels for numerous reasons including lack of ignition, successful fire suppression, or unfavorable weather. Moreover, the cost of underestimating burn probabilities among areas that do, in fact, burn (i.e., a false negative signal) is potentially greater than the cost of overestimating probabilities of areas that remain unburned (i.e., a false positive signal). We therefore scored predictions using the weighted log-loss, setting 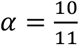 to weight predictions of burned pixels 10 times higher than predictions of unburned pixels. We selected as our final model the set of hyperparameters that minimized the global weighted log-loss across the tuning set.

#### 4. Pixel-level model evaluation

Samples from the 6 years withheld from model tuning were used to evaluate accuracy of probabilities from the tuned model at the 120 m pixel level. We report the log-loss as well as the more familiar classification accuracy metrics, kappa and AUC.

#### 5. Hindcasting

To evaluate model predictions at a practical spatial scale and provide information about the predictive utility that could be expected of future forecasts beyond what is provided by standard model performance metrics, we conducted a secondary validation based on hindcasts developed for each year from 1988–2019 (Fig. 2). As in our cross-validation model tuning procedure, hindcasts were constructed by withholding one year’s data at a time, fitting the model to samples drawn from all remaining 31 years, and then mapping predicted probabilities for the withheld year. These 32 hindcasts were then evaluated against actual burn footprints.

**Figure 2.**
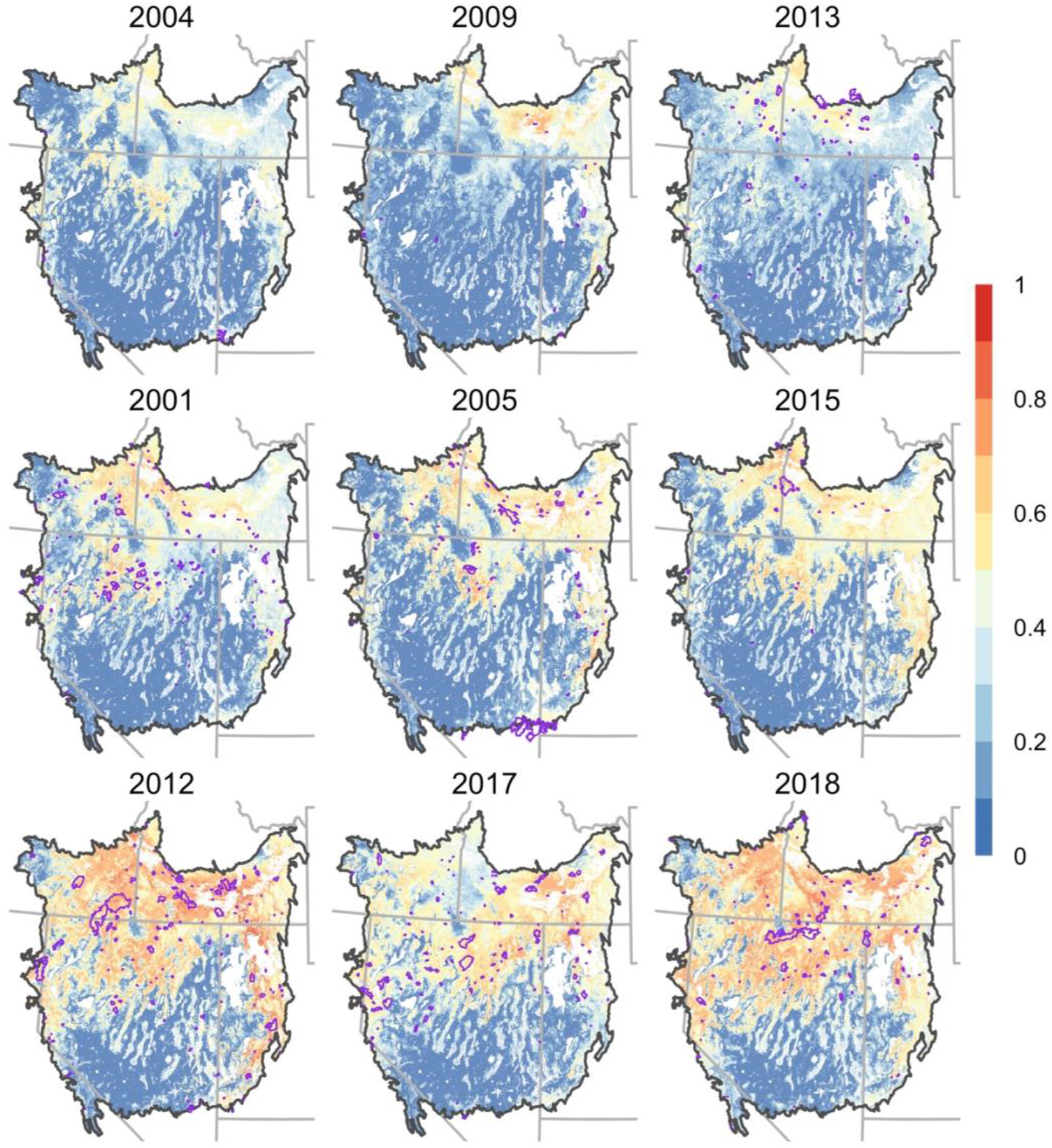
Hindcasts of fire probability in the Great Basin, USA, predicted by random forest models using antecedent cover and production of vegetation, multitemporal scale drought indices, and climate variables. Perimeters of wildfires >405 ha are depicted in purple. A leave-one-out approach was used to generate predictions for each year, with that year’s data withheld from model training. The nine most recent years with complete MTBS fire perimeter data are shown. An animation of the full time series of hindcasts, 1988–2019, is available online (https://rangelands.app/great-basin-fire/).

#### 6. Hindcast evaluation

We evaluated hindcasts by 1) converting continuous pixel-level estimates, 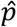, to ten decile bins, 2) overlaying the binary burned/unburned maps derived from the MTBS thematic burn severity dataset (see Data sources, above), and 3) calculating i) total area (roughly equal across bins) and ii) burned area in each bin. We then derived the proportion of the total area in each bin that burned. This proportion should increase monotonically from the lowest to highest bin if model fit is adequate. We also compared performance among forecast dates using a precision-recall curve based on the same aggregate decile bin data. Considering the lower bound of each bin sequentially as prediction thresholds, precision was approximated by the burned area for which 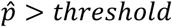 divided by the total area for which 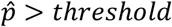. Recall was approximated by the burned area for which 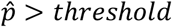 divided by the total burned area across bins. Higher precision for a given recall indicates higher skill in discriminating between pixels likely and unlikely to burn.

To evaluate correspondence between hindcasts and fire activity at the scale of the Great Basin over time, we aggregated predicted fire probability across the study area (mean) and calculated Pearson correlations between annual mean predicted fire probabilities and several annual metrics of fire season activity. These included total area burned in large wildfires, number of large wildfires, maximum fire size, and season length (number of days between ignition date of the first and last large wildfires). Total area burned was calculated using the MTBS thematic burn severity dataset to account for unburned inclusions within fire perimeters, while all other metrics were derived from the MTBS fire perimeter dataset. Highly skewed variables (total area burned, number of fires, max fire size) were either square-root (areas) or log (number of fires) transformed.

### Forecast date

To explore the trade-off between forecast lead time and accuracy, we tested forecasts made every 16 days from January 1 to May 25, aligning with the schedule of 16-day NDVI composites used to produce NPP estimates (Jones et al. 2021). For reference, between 1988 and 2019 the median ignition date of the first large wildfire in the Great Basin was March 30, and 95% of large wildfires started on or after June 12. Variables quantifying cumulative growing degree days and vegetation production in the current year (gdd, afgAGB_ytd, and pfgAGB_ytd) were omitted for January 1 forecasts. We repeated steps 1–6 for each forecast date, and report cross-validation metrics for each date.

### Variable importance and effects

We computed the permutation-based mean decrease in accuracy (hereafter permutation importance) and conditional importance to assess the relative importance of variables. Although permutation importance is an established method of ranking variables from RF models, it can be biased in the presence of highly correlated predictors (Strobl et al. 2007, Nicodemus and Malley 2009). The predictors we used were characterized by many high pairwise correlations. The conditional importance (Strobl et al. 2007, 2008) was developed as a less biased metric of variable importance under strong multicollinearity and other conditions that distort traditional variable importance metrics. We therefore calculated the conditional importance from conditional inference forests fit with the party package (version 1.3; Hothorn et al. 2006, Strobl et al. 2007, 2008) in R and compared importance rankings based on these two metrics. We also provide partial dependence plots to visualize the effects of individual predictors on fire probability. Partial dependence plots were produced for the 12 predictors ranking highest in conditional importance.

## Results

### Model tuning and evaluation

For all forecast dates, models constructed by building trees with a small subset of the training data sampled without replacement (sample fraction = 0.1) minimized the log-loss (Table 2). Cross-validation performance was less sensitive to *m*_*try*_, but values near random forest defaults 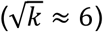 tended to perform well (Table 2). Pixel-level model performance on the withheld independent testing dataset was similar to performance in cross-validation, with threshold-based performance metrics (i.e., AUC, kappa) slightly improved, but log-loss slightly higher (Table 2).

**Table 2.** Model tuning and pixel-level evaluation of fuel-based rangeland fire probability models for the Great Basin using drought indices and concurrent year’s cumulative precipitation, growing degree days, and vegetation production at the indicated forecast date (DOY = day of year), and vegetation cover and annual production from the past 2 years. Performance in cross-validation and on independent testing data is shown for the optimal combination of random forest hyperparameters for each forecast date.

| Forecast<br>DOY | $m_{try}^1$ | Sample<br>fraction<sup>2</sup> | Cross-validation | | | Independent testing set | | |
| --- | --- | --- | --- | --- | --- | --- | --- | --- |
| | | | AUC | kappa | $L^*_{log}$ | AUC | kappa | $L^*_{log}$ |
| 1 | 5 | 0.1 | 0.851 | 0.524 | 0.27 | 0.868 | 0.552 | 0.33 |
| 17 | 4 | 0.1 | 0.849 | 0.525 | 0.27 | 0.870 | 0.522 | 0.34 |
| 33 | 4 | 0.1 | 0.853 | 0.521 | 0.27 | 0.868 | 0.565 | 0.32 |
| 49 | 6 | 0.1 | 0.847 | 0.510 | 0.27 | 0.871 | 0.575 | 0.32 |
| 65 | 4 | 0.1 | 0.851 | 0.521 | 0.27 | 0.872 | 0.570 | 0.33 |
| 81 | 6 | 0.1 | 0.856 | 0.535 | 0.26 | 0.873 | 0.582 | 0.32 |
| 97 | 5 | 0.1 | 0.846 | 0.518 | 0.27 | 0.866 | 0.548 | 0.34 |
| 113 | 5 | 0.1 | 0.841 | 0.483 | 0.28 | 0.863 | 0.545 | 0.33 |
| 129 | 5 | 0.1 | 0.851 | 0.518 | 0.27 | 0.870 | 0.554 | 0.34 |
| 145 | 5 | 0.1 | 0.855 | 0.525 | 0.27 | 0.862 | 0.538 | 0.34 |

<sup>1</sup> The RF hyperparameter $m_{try}$ controls the number of randomly selected predictors to consider at each split for building trees.
<sup>2</sup> Sample fraction refers to the fraction of training cases subsampled to train each tree.

### Forecast date

At the pixel level, accuracy was less sensitive to forecast date than expected (Table 2). In cross-validation, the log-loss was minimized on the March 22 (DOY 81) forecast date, but January 1 and May 25 (DOY 145) forecasts performed similarly (Table 2). We therefore selected 3 forecast dates—one early (January 1), one middle (March 22) and one late (May 25)—for evaluating hindcast performance and variable importance.

### Hindcast evaluation

Proportional area burned increased monotonically with increasing predicted fire probability for all three forecast dates (Fig. 3A). This contrasted with the static burn probability map (Short et al. 2020); although proportional burned area increased steadily through the 8^th^ decile bin, roughly equal proportions of the upper 3 decile bins burned (Fig. 3A).

**Fig. 3.**
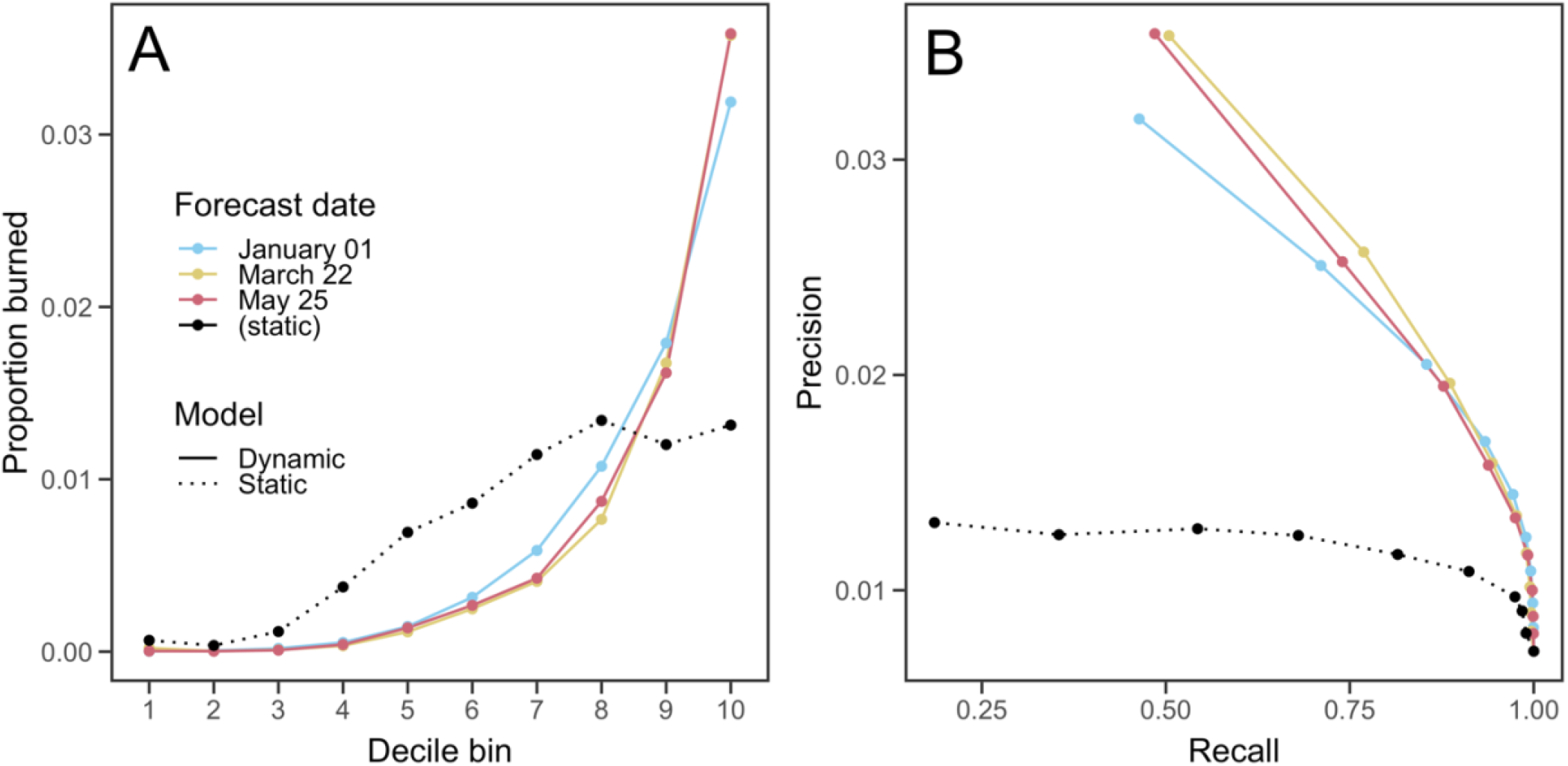
Validation of fuel-based rangeland fire probability hindcasts (“Dynamic”) using weather and vegetation production data through the indicated forecast date. Continuous predictions of large fire probability 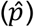 were divided into 10 equal-area bins. Panel A shows the proportion of the total mapped area in each bin that burned in large wildfires across years (1988–2019). Panel B shows the precision-recall tradeoff for each forecast date (see methods for calculations). Performance of a widely-used static burn probability map (“Static”) is shown for comparison (dashed black line).

Precision-recall curves revealed that hindcast skill increased only slightly from the earliest to the latest forecast date (Fig. 3B). Dynamic hindcasts consistently outperformed the static burn probabilities (Fig. 3B). The upper decile of probabilities from the January 1 hindcasts, for example, captured 43.8% of the total burned area, while the upper decile of probabilities from the static map captured only 18.5% of the total burned area. The upper 3 deciles of January 1 hindcasts captured approximately 88% of the cumulative burned footprint, compared to only 54% captured by the upper 3 deciles of static burn probabilities. 3.2% of the area in the upper decile from January 1 hindcasts burned, while only 1.3% of the upper decile of static burn probabilities burned.

Across years, mean predicted fire probability was strongly positively correlated (*ρ* > 0.6) with total burned area and number of large fires, and moderately positively correlated (0.6 > *ρ* > 0.4) with maximum fire size and fire season length (Fig. 4). These correlations varied little across the 3 examined forecast dates, although correlations were slightly stronger for the two earlier dates (January 1 and March 22).

**Figure 4.**
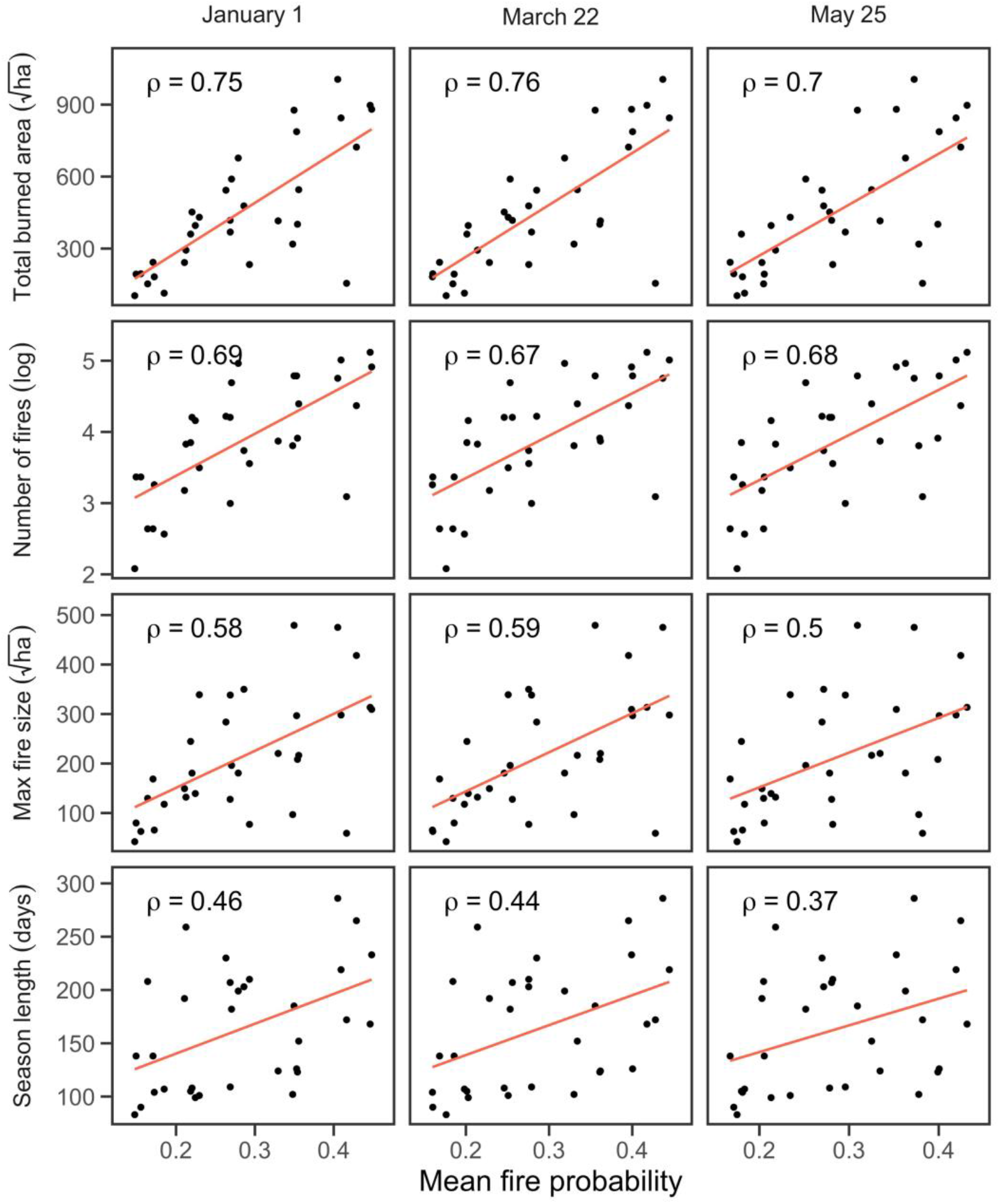
Metrics of annual wildfire activity in the Great Basin, USA, from 1988–2019 in relation to mean predicted fire probability. Mean fire probability was moderately to strongly correlated with the number of large wildfires, total area burned, and maximum fire size, and weakly correlated with fire season length. Mean predicted fire probability was calculated from January 1 (left column), March 22 (middle column) and May 25 (right column) forecast dates. Fitted lines are from least squares regressions.

### Variable importance

Permutation importance and conditional importance metrics agreed on the most important categories of variables (Fig. 5), including past years’ herbaceous vegetation cover and production, bare ground cover, and drought indices measured at 1- and 2-yr temporal scales. The highest-ranking vegetation variable was total herbaceous production in the previous year for both metrics, with the exception of conditional variable importance for the May 25 forecast. The 2-yr SPEI was also highly important, ranking in the top 3 variables for both metrics across forecast dates. Both metrics also agreed on the relatively low importance of shrub and litter cover, mean annual temperature (bio01), short-term drought indices (e.g., spei30d and eddi30d), and growing degree day accumulation (gdd). Lag-2 vegetation variables ranked lower in importance than lag-1 variables. A notable disagreement between permutation and conditional importance rankings was for tree cover; permutation importance placed tree cover as the lowest-ranked variable, while conditional importance ranked tree cover near the middle. However, because of the highly skewed distribution, even the conditional importance for tree cover was quite low relative to variables such as total herbaceous production and 2 yr SPEI.

**Figure 5.**
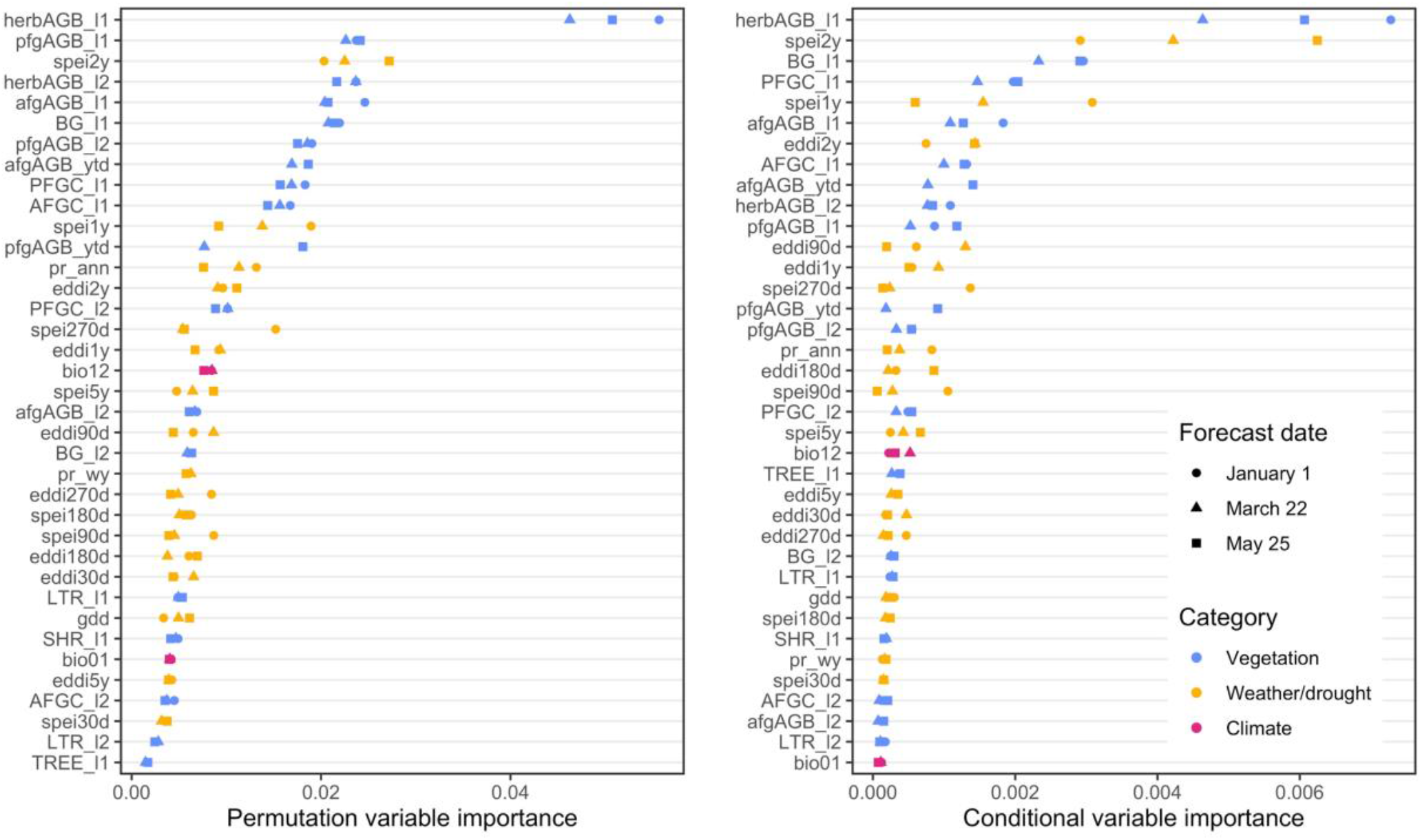
Importance of variables used to predict fire probability in the Great Basin, USA, 1988– 2019, corresponding to 3 forecast lead times (forecast date). The commonly-used permutation variable importance (left) yielded rankings that were generally similar to the conditional variable importance (right), which accounts for correlations among predictors. Antecedent production of herbaceous vegetation, long-term (1- to 2-year) drought indices, and bare ground were the most important predictors regardless of importance metric.

As anticipated, cumulative 16-day production of perennial and annual herbaceous vegetation (pfgAGB_ytd and afgAGB_ytd) were more important for forecasts made later in the spring (i.e., May 25) than those made earlier (i.e., March 22). In contrast, 1-yr SPEI was highly important for January 1 forecasts, but progressively diminished in importance for later forecast dates (Fig. 5).

## Discussion

Our analyses demonstrate how high-resolution, dynamic, remotely sensed vegetation data can be used to map fuels conducive to wildfires months in advance of the fire season in Great Basin rangelands. Annual hindcasts were predictive of both spatial and interannual patterns in fire activity over the last three decades, providing improved skill over static burn probabilities. Because the most important predictors were the previous year’s herbaceous vegetation production and long-term (1- to 2-yr scale) drought indices, accuracy was less sensitive to forecast lead time than anticipated. Thus, forecasts providing timely and accurate information about the spatial distribution of fuels and the potential severity of the upcoming fire season can be made available in early January, potentially expanding opportunities for advanced planning and resource allocation for wildfire preparedness.

The strong correlations between average predicted fire probability and annual metrics of fire activity (number of large fires, total area burned, etc.; Fig. 4) speak to the extent to which fire in Great Basin rangelands is controlled by antecedent conditions related to the growth of herbaceous vegetation as opposed to fuel moisture or fire weather; i.e., it is a fuel-limited system. This finding is consistent with past research (e.g., Knapp 1998, Abatzoglou and Kolden 2013, Pilliod et al. 2017), and suggests information about fuel quantity (i.e., biomass) is equally or more important than information about fuel quality (i.e., moisture content) in sagebrush rangelands.

A role of fuel moisture is, however, suggested by effects of shorter-term drought indices (Fig. S2, S3). For example, the opposite effects of EDDI at longer (e.g., 2 yr) and shorter (e.g., 90 d) temporal scales indicates fire probability is maximized when a long-term moisture surplus (i.e., SPEI > 0 or EDDI < 0 at 1-2 yr scales) is followed by a dry winter/spring (Fig. S2). Among later forecast dates, however, the effect of current year’s growth of fine fuels (afgAGB_ytd and pfgAGB_ytd) becomes increasingly important (Fig. 5, S3). These opposing effects of spring moisture on fire probability may help explain the weak correlation between spring conditions and annual fire activity in rangelands of the Great Basin (Fig. S4).

Evidence of weather’s influence during the fire season also appears in a few notable outliers in the regression of burned area on mean fire probability. These include years characterized by particularly wet late spring and summer conditions in which the burned area was well below expectations (e.g., 1997; Fig. S5) and years of above average summer temperatures in which the burned area exceeded expectations (e.g., 2007; Fig. S5). Weather is also highly influential in determining the intra-annual timing of fire activity and outcomes of individual fires, and therefore fire danger indices based on vegetation flammability and atmospheric conditions will continue to play a central role in short-term preparedness and fire suppression.

Herbaceous fuels—forbs and grasses—were the most influential predictors of fire probability, whereas woody fuels (trees and shrubs) were relatively unimportant (Fig. 5). Inference regarding the effects of tree cover was limited, however, by the way we defined our study area; most regions exceeding approximately 5% tree cover were excluded. Although the presence and abundance of woody vegetation increases flame lengths, fire intensity, and residence time with potential negative implications for suppression and post-fire recovery (Strand et al. 2013, Boyd et al. 2015, Weiner et al. 2016), the probability of burning *per se* is controlled primarily by characteristics of the herbaceous understory. To the extent that certain woody species (e.g., western juniper; Miller et al. 2000) can reduce herbaceous cover, higher woody fuel loads may in fact be associated with reduced incidence of fire (Miller and Tausch, 2001). Moreover, certain treatments to reduce woody vegetation can increase herbaceous fuels, e.g., through soil disturbance or release from competition (Pyke, 2011, Pyke et al. 2014). An effective fuel management program must consider the distinct roles of herbaceous and woody fuels and the likely effects of management interventions on both fuel types. If reducing the incidence of large wildfires is the objective, fuel management should emphasize the reduction of herbaceous fuel loads. Where fire suppression effectiveness and safety are primary concerns, woody fuel treatments may be appropriate.

Although intense interest has attended the role of exotic annual grasses in fueling large fires in the Great Basin and throughout the western US (Balch et al. 2013, Davies and Nafus 2013, Fusco et al. 2019), perennial grasses and forbs were equally important as predictors of burning (Fig. 5). Total annual herbaceous production in the previous year was the highest in overall importance across metrics and forecast dates (with the exception of conditional importance for May 25 forecasts). This suggests the composition of herbaceous fine fuels may be less critical than the quantity for predicting occurrence of fire. While there is ample evidence exotic annual grasses have reduced fire return intervals in western US rangelands (Balch et al. 2013), ecoregional-scale interannual variation in fire activity remains strongly controlled by weather-driven variation in productivity of perennial herbaceous vegetation (Pilliod et al. 2017).

Long-term trends in fire activity, on the other hand, appear to be driven predominantly by increasing cover and production of annuals. A trend of increasing annual burned area has recently emerged in the Great Basin (Fig. S6), with the 12 largest fires in the MTBS record within our study area having occurred since 2000 (of those, 8 occurred since 2010). This mirrors increases in fire activity occurring across ecosystems of the western US—a trend which has been attributed in part to the emerging influence of anthropogenic climate change on fire weather and fire season length (Jolly et al. 2015, Abatzoglou and Williams 2016, Abatzoglou et al. 2019, Bowman et al. 2020). Among the shrublands and grasslands of the Great Basin, however, the upturn appears to be driven largely by increasing fine fuels—in particular, exotic annual grasses. While perennial forb and grass production has remained stationary to slightly decreasing, cover and production of annuals has increased rapidly (Pastick et al. 2021, Smith et al. 2022), with an attendant rise in average fire probability (Fig. S6). This trend in fire probability, in turn, appears to largely explain the increase in burned area (Fig. 4; Fig. S6), with little indication that variation in burned area has become decoupled from variation in fuel quantity. We verified this with a *post-hoc* analysis in which we fit a linear regression relating total annual burned area 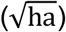 to mean herbaceous production (kg·ha^−1^) in the previous year (the most important predictor from our models) at the scale of the Great Basin. The fitted regression had an R^2^ of 0.48, and residuals exhibited no temporal trend (Mann-Kendal *Z* = -0.34, p = 0.73). This implicates increasing fine fuels, rather than more extreme fire weather or longer fire seasons, as the primary factor responsible for increasing fire activity in the Great Basin (also see Balch et al. 2013).

### Implications

The ascendancy of fine fuels as drivers of interannual and spatial patterns in wildfire activity in Great Basin rangelands underscores the potential efficacy of fuel management for mitigating wildfire hazard. Alongside ignitions, fuels are among the only factors that can be pre-emptively addressed by land managers. Although fine fuels exert strong control on fire in these ecosystems, other factors such as extreme fire weather and national-scale demand on resources still play an important role in the outcomes of individual fires and fire seasons. The combination of rising production by invasive annual grasses, weather-driven peaks in perennial forbs and grass production, and synchronous extreme fire weather are likely to increasingly challenge the capacity of fire suppression resources and cause wildfire disasters. However, long-lead, fuels-based spatial forecasts such as those developed here may help managers anticipate and mitigate such events. Managers monitor a suite of indicators of fire danger, including drought indices, snowpack, long-term weather outlooks, and vegetation moisture data as the fire season approaches. Because these indicators vary widely in spatial and temporal grain and extent, their integration into a coherent picture of the regional distribution of fuels leading into the fire season remains challenging, particularly at sub-ecoregional scales. A major contribution of the framework we present here is the integration of many of the indicators already widely used by managers into a single predictive metric applicable across a broad range of spatial scales, from the pixel to the ecoregion (Fig. 6). Used alongside short-term fire danger indices and other tools, these forecasts may help managers more accurately predict and prepare for where and when ignitions are likely to result in expensive, dangerous, and ecologically damaging large wildfires.

**Figure 6.**
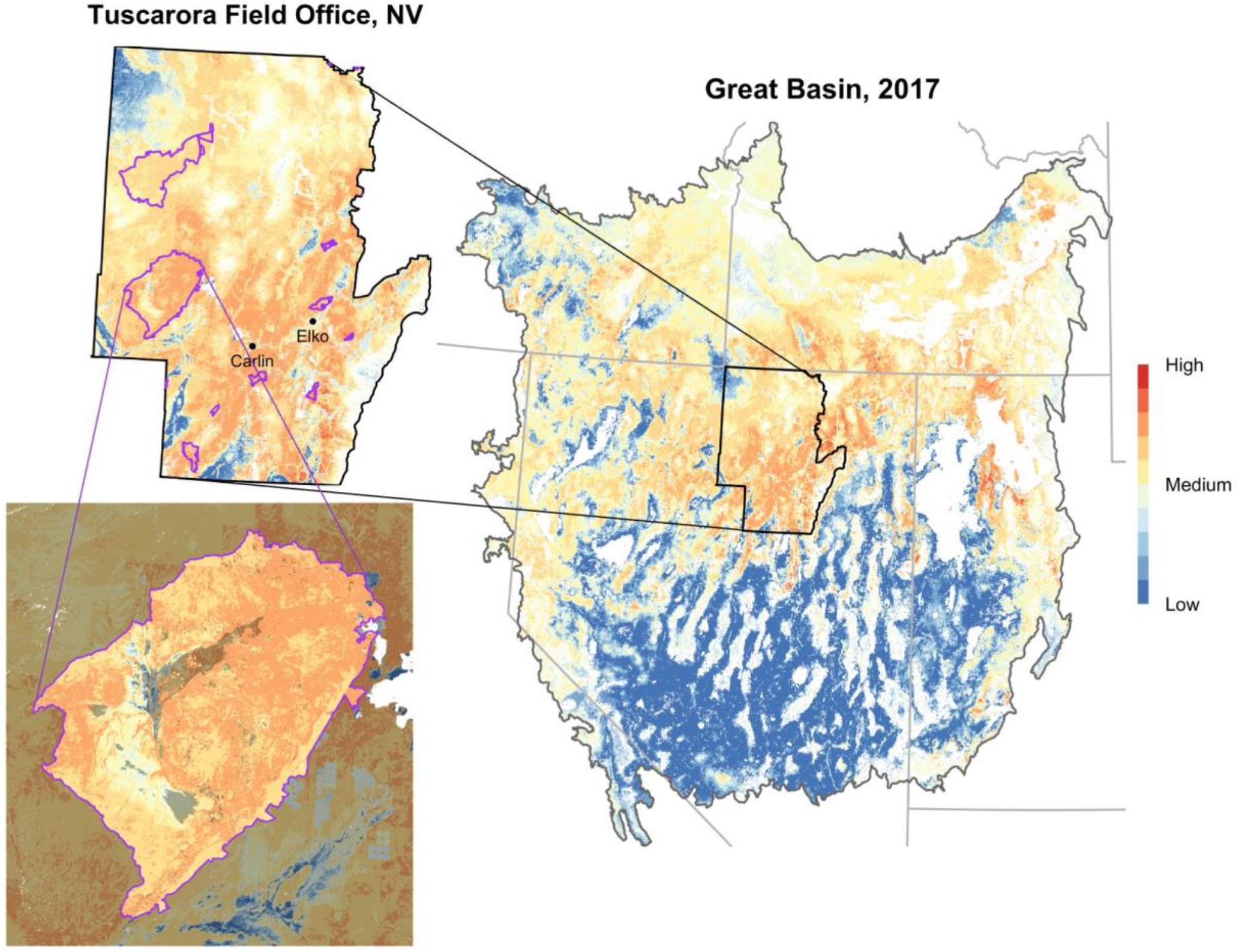
2017 fire probability in the Great Basin, USA (March 22 forecast date). Insets highlight the applicability across a range of spatial scales. Used in conjunction with fire danger indices, other fire forecasts, and expert knowledge, seasonal fire probability maps may help managers at multiple levels prepare for where and when ignitions are likely to result in large and damaging wildfires. Perimeters of large (>405 ha) wildfires that occurred in 2017 are depicted in purple within insets. The lower left inset shows the 2017 Rooster Comb fire, with darker shading indicating unburned areas.

## Acknowlegements

We thank Casey O’Connor and Mike Pagoaga for providing insightful feedback from a fire management perspective, and Stella Copeland, Dustin Johnson, and two anonymous reviewers for their constructive reviews of the manuscript. This work was supported by the U.S. Department of Agriculture, Agricultural Research Service. The findings and conclusions in the publication are those of the authors and should not be construed to represent any official USDA or U.S. Government determination or policy. USDA is an equal opportunity provider and employer.

