## Supplementary material for "Where there’s smoke, there’s fuel: dynamic vegetation data improve predictions of wildfire hazard in the Great Basin"

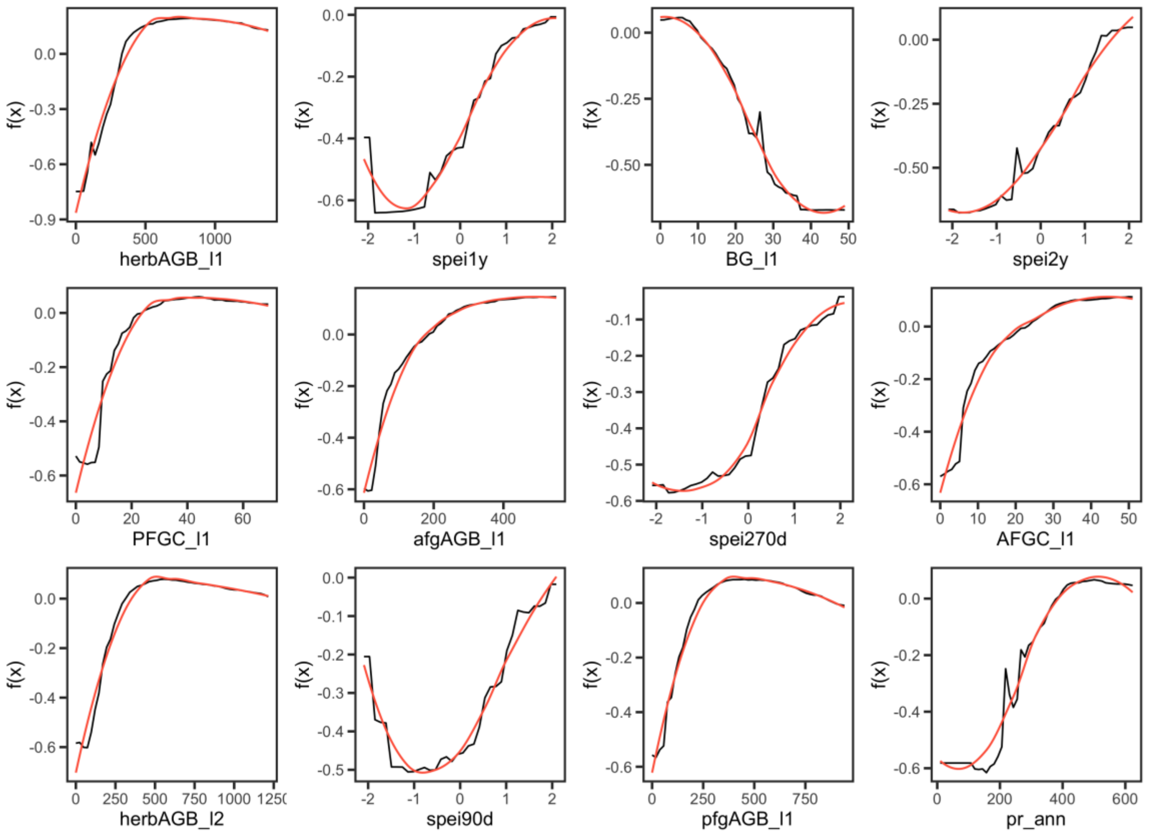


Figure S1. Partial dependence plots depicting the effects of the 12 top-ranked predictors (by conditional variable importance) on the relative probability of burning in a large (>405 ha) wildfire in the Great Basin for January 1 forecasts. Black lines show the partial dependence function $f(x)$ (centered logit of fire probability), with the general shape of the relationship highlighted with a LOESS smoother (span = 0.75; red lines).

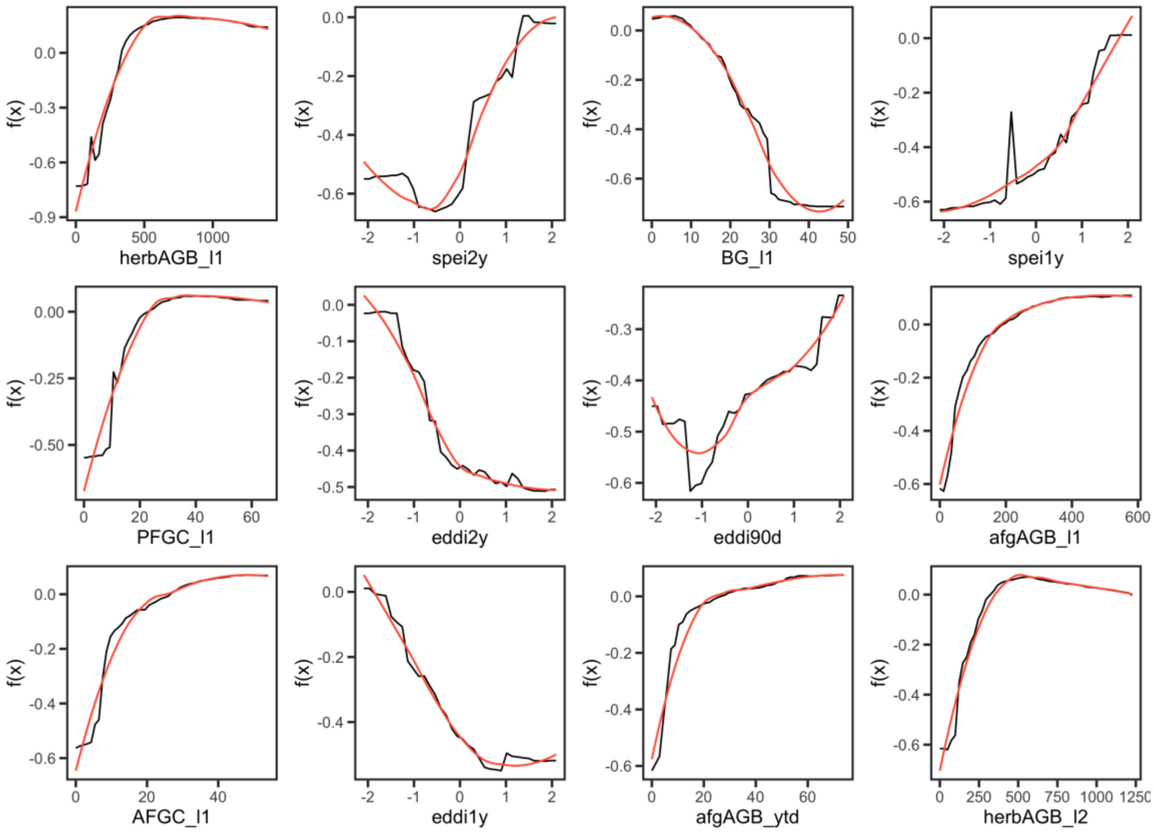


Figure S2. Partial dependence plots depicting the effects of the 12 top-ranked predictors (by conditional variable importance) on the relative probability of burning in a large (>405 ha) wildfire in the Great Basin for March 22 forecasts. Black lines show the partial dependence function $f(x)$ (centered logit of fire probability), with the general shape of the relationship highlighted with a LOESS smoother (span = 0.75; red lines).

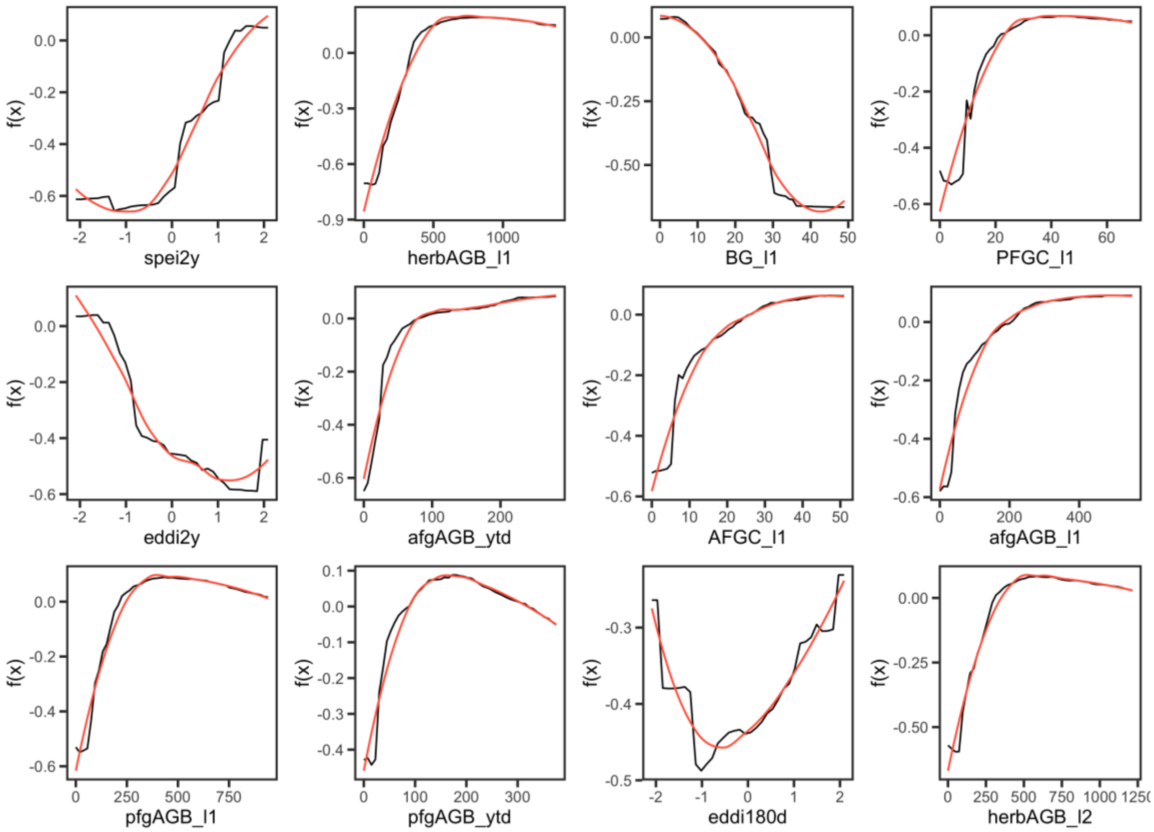


Figure S3. Partial dependence plots depicting the effects of the 12 top-ranked predictors (by conditional variable importance) on the relative probability of burning in a large (>405 ha) wildfire in the Great Basin for May 25 forecasts. Black lines show the partial dependence function $f(x)$ (centered logit of fire probability), with the general shape of the relationship highlighted with a LOESS smoother (span = 0.75; red lines).

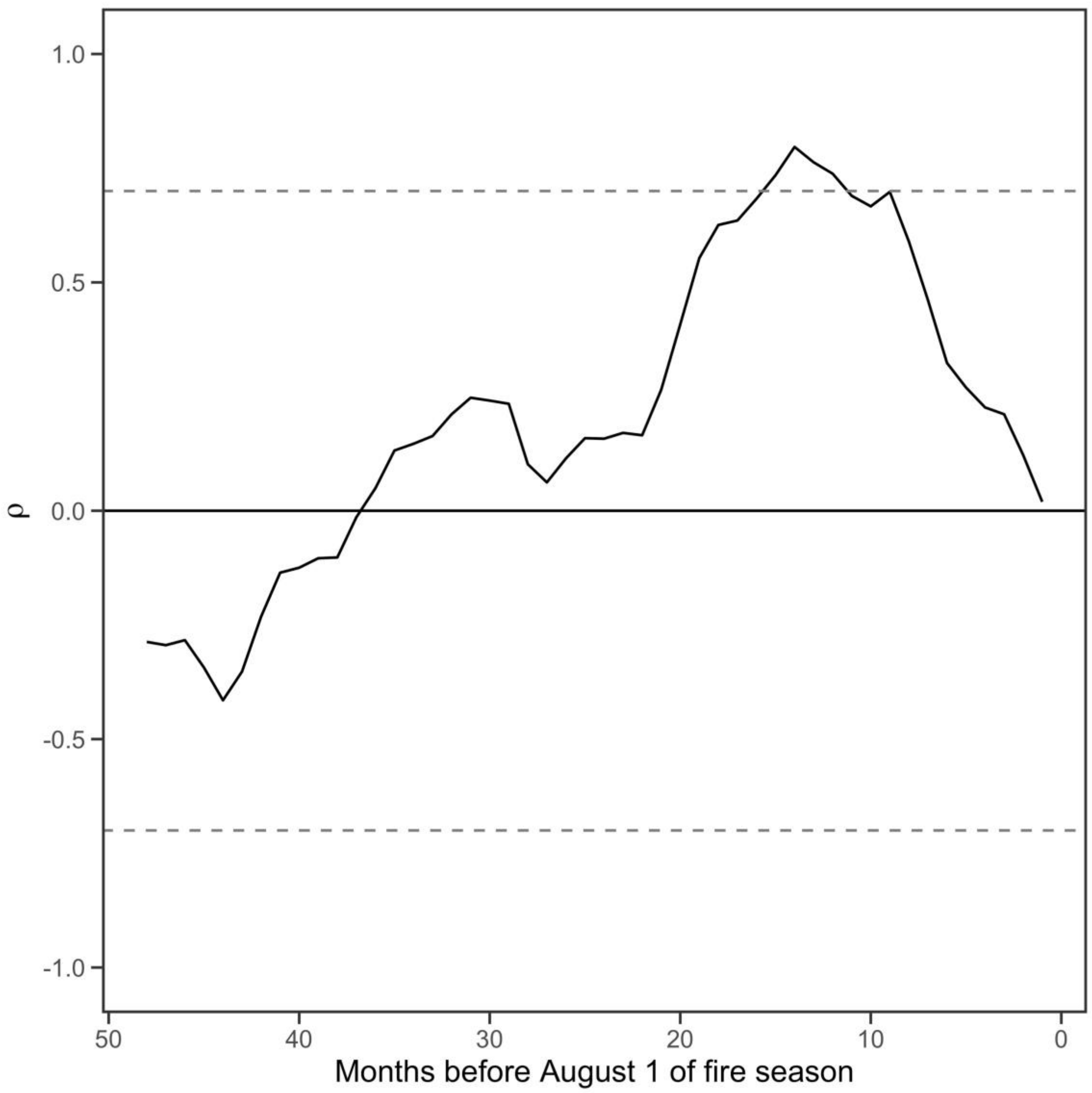


Figure S4. Pearson correlation between the Palmer Drought Severity Index (PDSI) and total area burned in large (>405 ha) wildfires across the Great Basin, 1986–2019, for lags of 0 to 48 months from August 1 of the fire season. Dashed lines indicate correlations of -0.7 and 0.7. Burned area is most strongly correlated ($\rho$ = 0.797) with the PDSI measured at the end of May in the previous year, demonstrating the importance of antecedent growth of fine fuels in controlling fire in these ecoregions.

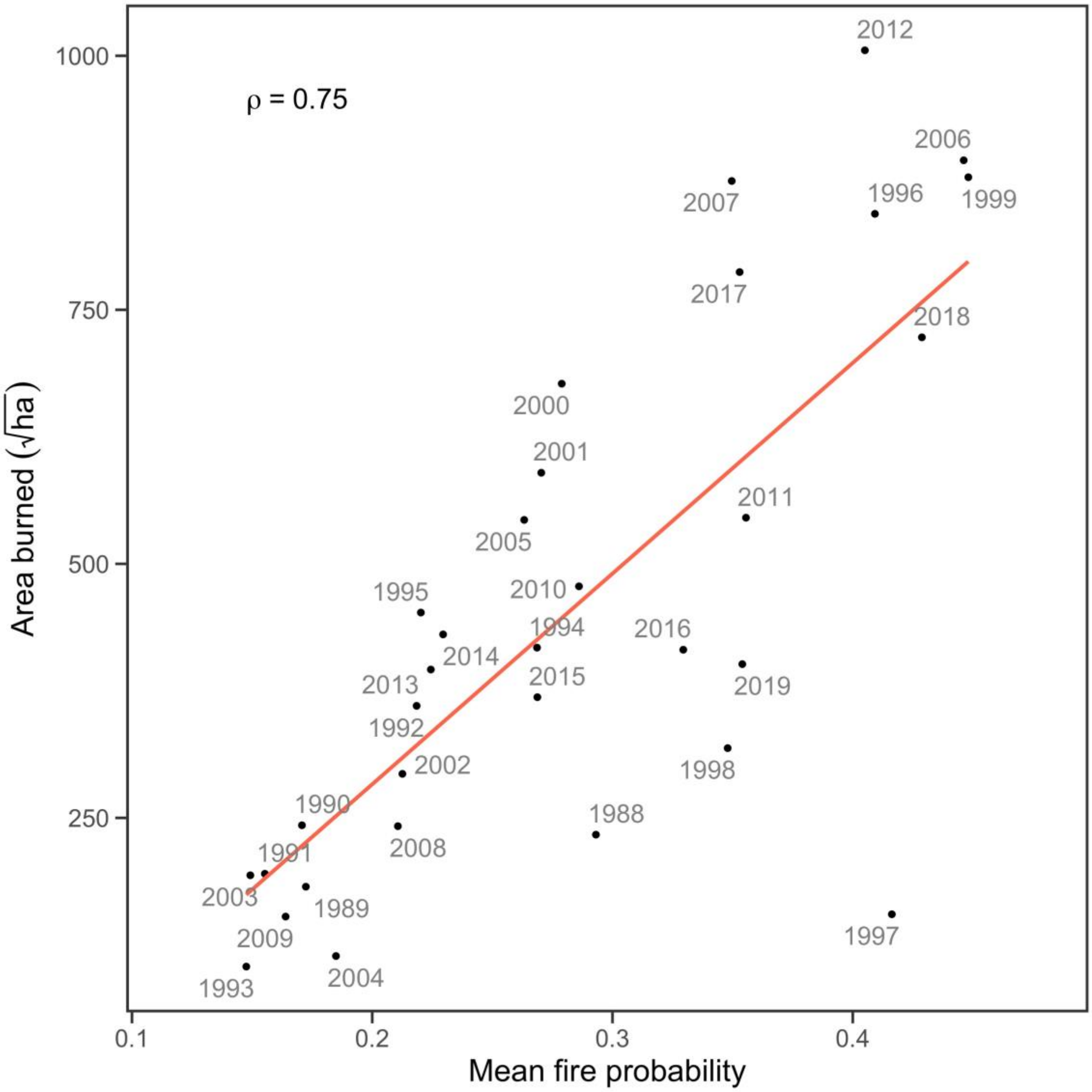


Figure S5. Relationship between mean predicted fire probability and total area burned in large (>405 ha) wildfires in Great Basin rangelands, 1988–2019. Mean fire probability was calculated from January 1 hindcasts (same data as shown in top-left panel in Fig. 4, but with years labelled).

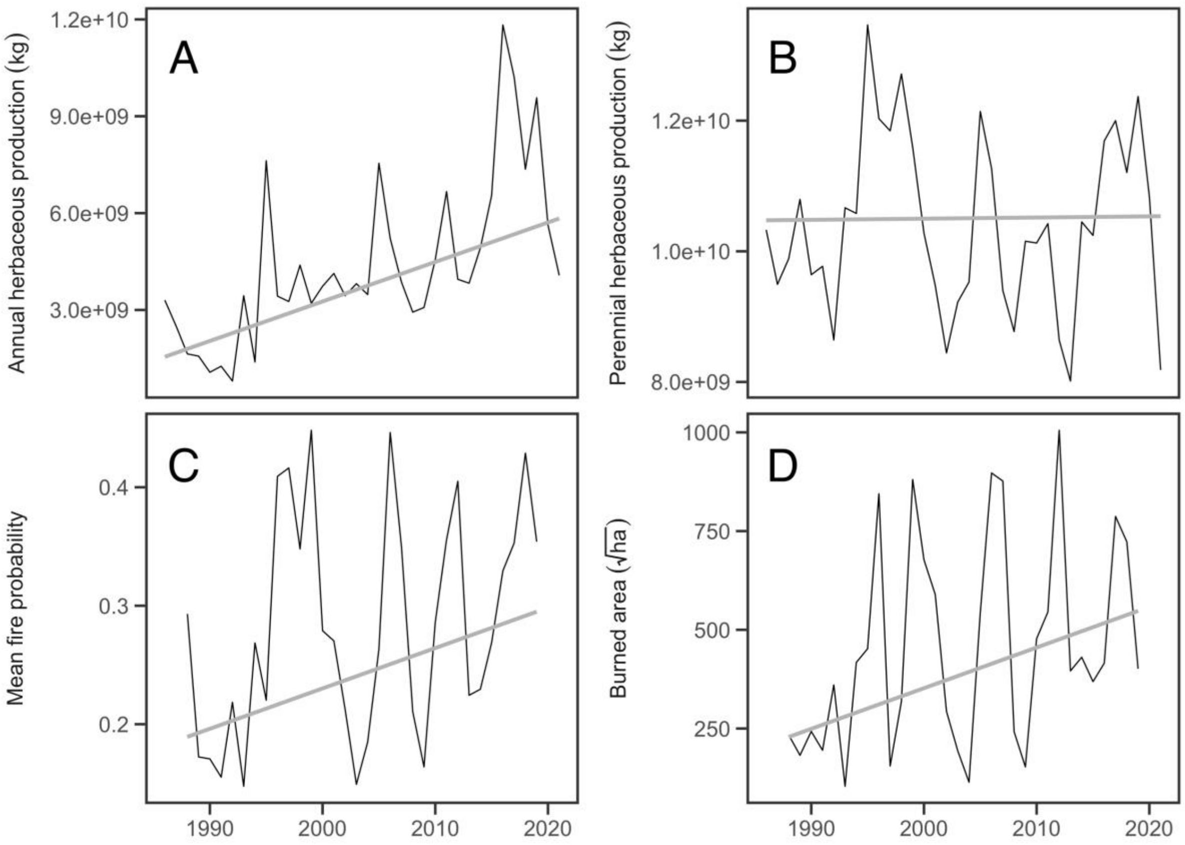


Fig. S6. Trends in total annual (A) and perennial (B) herbaceous aboveground biomass production across rangelands of the Great Basin, 1986–2021. Production of annuals has steadily increased while production of perennials showed no long-term trend. Increases in predicted fire probability (C; January 1 hindcasts), and area burned in large (>405 ha) wildfires (D) from 1988–2019 parallel the increase in production of annuals. Fitted lines are Thiel-Sen slope estimates.
